# Genetic Mouse Models of Autism Spectrum Disorder Present Subtle Heterogenous Cardiac Abnormalities

**DOI:** 10.1101/2021.10.19.465007

**Authors:** Stephania Assimopoulos, Christopher Hammill, Darren J. Fernandes, Tara Leigh Spencer Noakes, Yu-Qing Zhou, Lauryl M. J. Nutter, Jacob Ellegood, Evdokia Anagnostou, John G. Sled, Jason P. Lerch

**Affiliations:** Mouse Imaging Centre, Hospital for Sick Children, Toronto, Ontario, Canada; The Hospital for Sick Children, Toronto, Ontario, Canada; Department of Medical Biophysics, University of Toronto, Toronto, Ontario, Canada; The Centre for Phenogenomics, Toronto, Ontario, Canada; Holland Bloorview Kids Rehabilitation Hospital, Toronto, Ontario Canada; Wellcome Centre for Integrative Neuroimaging, The University of Oxford, Oxford, UK

**Keywords:** autism, cardiac phenotype/cardiology, comorbidities, genetic mouse models, phenotyping, ultrasound biomicroscopy

## Abstract

**Background:** Autism Spectrum Disorder (ASD) and Congenital Heart Disease (CHD) are strongly linked on a functional and genetic level. Most work has been focused on neurodevelopmental abnormalities in CHD. Conversely, cardiac abnormalities in ASD have been less studied. In this work we investigate the prevalence of cardiac comorbidities relative to genetic contributors of ASD.

**Methods:** Using high frequency ultrasound imaging, we screened 9 mouse models with ASD-related genetic alterations (*Arid1b*^*(+/-)*^, *Chd8*^*(+/-)*^, 16p11.2 (deletion), *Sgsh*^*(+/-)*^, *Sgsh*^*(-/-)*^, *Shank3 Δexon 4-9*^*(+/-)*^, *Shank3 Δexon 4-9*^*(-/-)*^, *Fmr1*^*(-/-)*^, *Vps13b*^*(+/-)*^), and pooled wild-type littermates (WT). Using a standardised imaging protocol, we measured heart rate (HR), aorta diameter (AoD), thickness and thickening of the left-ventricular (LV) anterior and posterior walls, LV chamber diameter, fractional shortening, stroke volume and cardiac output, Peak E and A velocity ratio of mitral inflow, Velocity Time Integral (VTI) through the ascending aorta.

**Results:** Mutant groups presented small-scale alterations in cardiac structure and function compared to WTs. A greater number of significant differences was observed among mutant groups than between mutant groups and WTs. Mutant groups differed primarily in measures of structure (LV chamber diameter and anterior wall thickness, HR, AoD). When compared to WTs, they differed in both structure and function (LV anterior wall thickness and thickening, chamber diameter and fractional shortening, HR). The mutant groups with most differences to WTs were 16p11.2 (deletion), *Fmrl*^*(-/-)*^, *Arid1b*^*(+/-)*^. Among mutant groups, the groups differing most from others were 16p11.2 (deletion), *Sgsh*^*(+/-)*^, *Fmr1*^*(-/-)*^. Our results broadly recapitulate the associated clinical findings.

**Limitations:** Various genetically driven cardiac abnormalities occur early in life, so repeating this work in non-adult mice may be valuable. To identify possible sex differences, we must extend this work to female mice. The downsampling procedure used (total correlation calculation) must be verified. Only indirect comparison between our results and clinical literature is possible due to differing study designs.

**Conclusions:** The characteristic heterogeneity of ASD was recapitulated in the observed cardiac phenotype. The type of measures (morphological, functional) mutant groups differ in can highlight common underlying mechanisms. Clinically, knowledge of cardiac abnormalities in ASD can be essential as even non-lethal cardiac abnormalities can impact normal development.

## Background

Autism Spectrum Disorder (ASD) and Congenital Heart Disease (CHD) have been found to be strongly linked both on a functional (1–3) and a genetic level (4–6). ASD is a neurodevelopmental disorder (NDD) with an occurrence rate >1%, highly heterogeneous in etiology and phenotype (7,8). It is primarily associated with behavioural deficits, but also has associated comorbidities, amongst which cardiac are common (9). CHD is the most common birth defect (10) present in 0.8%-1.2% of all live births worldwide (11). It accounts for about 3% of infant deaths and, of those, 46% are due to congenital malformations (12). Neurodevelopmental differences, including language, motor, cognitive and social deficits, occur in 10% of children with congenital heart disease and in 50% of those with severe congenital heart disease (13).

The primary focus of the investigation in the NDD-CHD association, has been to identify the frequency of neurodevelopmental differences in various CHD cases. There are only a few studies on the ASD-CHD association, and they report an increased risk of ASD in CHD (6,14). In CHD, brain development is found to be atypical on MRI studies, where newborns with CHD were found to have smaller for gestational age total brain volume, reduced metabolism and delayed cortical development and folding compared to controls (following age and weight adjustment) (1,15–18), both prior to and following the necessary corrective cardiac surgeries (1,18). Moreover, known risk factors can only account for 30% of the neurodevelopmental outcome following such interventions, suggesting there are other genetic and epigenetic factors contributing to the outcomes (19). Reinforcing the association between cardiac and brain development on the genetic level, a large study performing exome sequencing on 1,213 congenital heart disease parent-offspring trios identified 69 genes containing damaging *de novo* mutations known to be associated with both ASD and NDD, all of which are in the top quartile of both developmental heart and brain expression (far more than expected by chance), thus revealing a shared genetic contribution to congenital heart disease and NDD (4).

Contrary to the aforementioned work on NDDs in CHD, the investigation of cardiac abnormalities in ASD, whether specifically CHD or not, has been more limited. The incidence rate of cardiac abnormalities reported in ASD varies in the literature depending on the cardiac abnormalities considered and the age of the subjects. In an adult U.S.A. population sample, Croen et al. (2015) considered three categories of cardiovascular diseases (dyslipidemia, hypertension and all other) and reported a joint prevalence of 36.96% and an odds ratio of 2.54, with hypertension being the most prevalent amongst the three groups (20). Similarly, in an adult French population subset, Miot et al. (2019) considered five categories of which three were significant (heart failure, orthostatic hypotension, peripheral vascular disease) and reported a joint prevalence of 15.37%, while separately considering hypertension with a prevalence of 13.56% (21). A population study in Western Australia assessing the occurrence rate of birth defects diagnosed before the age of 6 in children with ASD, reported a 1.3% incidence rate with a 1.2 odds ratio for cardiovascular system defects (22).

ASD is associated with a vast spectrum of cardiac abnormalities, including morphological alterations, cardiac dysrhythmias (ventricular flutter, fibrillation, and premature beats) and dysregulated resting autonomic activity. Autonomic dysregulation is one of the main abnormalities observed in syndromic cases of ASD (5,23). Specifically, cardiac dysrhythmias and various morphological defects are most prominent in chromosomal disorders of 22q11 deletion syndrome (DiGeorge, velocardiofacial syndrome) (24,25), in which the co-existence of a cardiac abnormality has been found to also result in reduced cortical and hippocampal volume (26). Additionally, cardiac arrythmias are commonly seen in patients with Timothy syndrome (*CACNA1C* mutation) (27,28). Similarly, CHARGE syndrome, a genetic disorder manifested through a non-random association of congenital anomalies (29), has a known associated cardiac dysfunction (30–32). Related investigations have been conducted for other NDDs, such as Trisomy21 (Down Syndrome) and Turner Syndrome, in which congenital heart disease is present in 50% and 30% respectively (23,33), as well as in Rett syndrome (34). Dysregulation in resting autonomic activity, or equivalently vagus nerve activity, is also observed in non-syndromic cases of ASD. Specifically, sympathetic overarousal, parasympathetic underactivity or atypical interaction between the two have been observed (35–37). Such dysregulation has been seen in both children and adults with ASD and is mainly manifested as reduced resting-state heart rate (HR) variability, slowed respiratory sinus arrythmia (RSA) and/or increased HR (36–38). Dysregulation in autonomic activity/vagus nerve function can generally lead to various cardiovascular complications thus warranting further investigation. Reports of morphological cardiac abnormalities have been sparse and mainly reported in syndromes associated with ASD, such as Tuberous Sclerosis (*TSC1* or *TSC2*) and Fragile X syndrome (5,23,39,40). Finally, accumulating evidence suggests that severe cardiac abnormalities do not simply co-occur with NDDs but may also contribute to the observed neurodevelopmental abnormalities (1,19,25), thus further highlighting the need for deeper investigation of cardiac abnormalities in ASD.

It is apparent that cardiac comorbidities in ASD may arise as a result of various, potentially coinciding, factors. In this work we investigate the prevalence of cardiac comorbidities relative to prominent genetic contributors of ASD. Specifically, using high frequency ultrasound imaging, we screen a set of ASD-related genetic mouse models with a standardised protocol to assess the incidence of cardiac abnormalities.

This study is performed in mice to allow for tightly controlled genetics, environment and experimental procedures. Studies in mouse models of ASD have been useful in determining the impact of genomic variation on brain differences across species (41). In addition, the development of the healthy mouse heart is comparable to that of humans and various mouse models of CHD have been reported to adequately replicate the corresponding human condition (42), suggesting that mouse models may be well suited for this endeavour.

## Methods

### Animals

Adult (60 days old +/- 1 day) male mice from 9 mouse models of 7 ASD-related genes (Table 1) were used in this study. The abnormalities and their effects, resulting from the underlying genetic mutations, can emerge at varying timepoints in development and potentially become more pronounced with time. Thus, by phenotyping our mice at an adult age, we aim to target potentially more stable and more pronounced phenotypes. Both heterozygous (+/-) and homozygous (-/-) mutant mice were assessed where possible. Our control group comprised of 2 wild-type (+/+) littermates from each model (with the exception of *Shank3* where we had 3), pooled across all models. The models for *Arid1b, Chd8, Vps13b*, and *Sgsh*, were produced at The Centre for Phenogenomics and provided by the Canadian Mutant Mouse Repository (CMMR, Toronto ON). They were all created on a C57BL/6NCrl background (Charles River Laboratories, strain code 027). The mouse models *Fmr1* (Jackson Laboratory #003025) (43), 16p11.2 deletion (Jackson Laboratory #013128) (44) and *Shank3 (Δexon 4-9)* (Jackson Laboratory #017890) (45) were obtained from Jackson Laboratory and were backcrossed at least seven generations to C57BL/6NCrl (Charles River Laboratories, strain code 027). All procedures involving animals were performed in compliance with the Animals for Research Act of Ontario and the Guidelines of the Canadian Council on Animal Care. The Centre for Phenogenomics (TCP) Animal Care Committee reviewed and approved all procedures conducted on animals at TCP. The number of mice comprising each group is also listed in Table 1 and was determined following a power analysis with significance level α=0.05 and power β= 0.85.

**Table 1:** Mouse models used in the study given by their common name and official mouseline name. Genotype and number of mice used per model also listed.

| Mouse Model – Common Name | Mouseline | Genotype | Number of Mice |
| --- | --- | --- | --- |
| Sgsh-Het | C57BL/6NCrI-Sgsh<sup>em3(IMPC)Tcp</sup>/Cmmr | (+/-) | 20 |
| Sgsh-Hom | C57BL/6NCrI-Sgsh<sup>em3(IMPC)Tcp</sup>/Cmmr | (-/-) | 20 |
| Fmr1 | B6:129P2-Fmr1/n | (-/Y) | 20 |
| Arid1b | C57BL/6NCrI-Arid1b<sup>em1(IMPC)Tcp</sup>/Cmmr | (+/-) | 20 |
| Shank3-Het (Δexon 4-9) | B6(Cg)-Shank3<sup>tm1.2Bux</sup>/n | (+/-) | 20 |
| Shank3-Hom (Δexon 4-9) | B6(Cg)-Shank3<sup>tm1.2Bux</sup>/n | (-/-) | 20 |
| 16p11.2 deletion | B6.129S-Del(7Slx1b-Sept1)4Aam/n | (df/+) | 20 |
| Chd8 | C57BL/6J-Chd8<sup>em1Tcp</sup> | (+/-) | 20 |
| Vps13b | C57BL/6NCrI-Vps13b<sup>em1(IMPC)Tcp</sup>/Cmmr | (+/-) | 20 |
| Wild-Type (WT) | - | (+/+) | 21 |

### Animal Preparation

Mice were prepared as described by Zhou et al. (46). They were anesthetized using isoflurane (induced at 5% in medical oxygen, and then maintained at 1.5% through a face mask). Mice were positioned supine with four paws taped to electrodes on a pre-warmed platform for ECG recording and heart rate monitoring. Mouse body temperature was monitored by rectal thermometer (Indus Instruments, Houston, TX) and maintained around 36-37°C by a heated platform. Mouse hair on the whole chest was cleanly removed using hair-removal cream (Nair, Carter-Horner, Mississauga, Ontario). Finally, a prewarmed ultrasound gel (Aquasonic 100; Parker Laboratories, Orang, NJ) was used as a coupling medium between the transducer and mouse body for image acquisition.

### Imaging

A high frequency ultrasound imaging system (Vevo 2100, FUJIFILM VisualSonics Inc., Toronto) with a 30MHz linear array transducer was used for assessing cardiac structure, function and hemodynamics, mainly following an imaging protocol published previously (46,47). Unbiased screening of all models was performed using a standardised battery of measures utilizing four conventional function modalities of ultrasound imaging for a full cardiac assessment. Table 2 lists the imaging modalities, approaches/sections and measurements of interest. With the exception of HR, values for all metrics were obtained by averaging the measurements from three cardiac cycles in the same ultrasound recording.

**Table 2:**
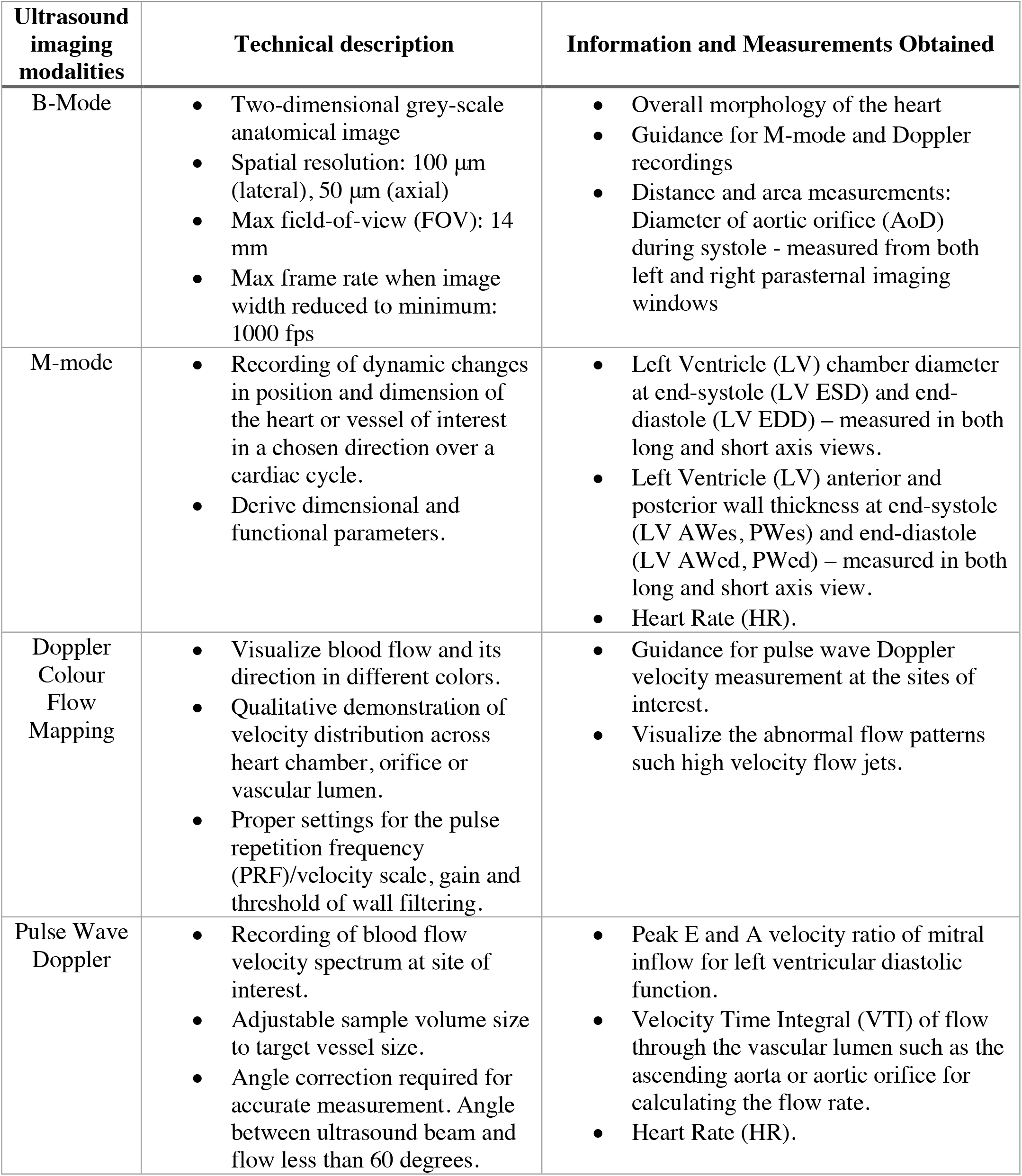
Four ultrasound imaging modalities used for full cardiac assessment, their technical description and the measurements obtained with each.

| **Ultrasound imaging modalities** | **Technical description** | **Information and Measurements Obtained** |
| --- | --- | --- |
| B-Mode | Two-dimensional grey-scale anatomical image<br>Spatial resolution: 100 $\mu\text{m}$ (lateral), 50 $\mu\text{m}$ (axial)<br>Max field-of-view (FOV): 14 mm<br>Max frame rate when image width reduced to minimum: 1000 fps | Overall morphology of the heart<br>Guidance for M-mode and Doppler recordings<br>Distance and area measurements: Diameter of aortic orifice (AoD) during systole - measured from both left and right parasternal imaging windows |
| M-mode | Recording of dynamic changes in position and dimension of the heart or vessel of interest in a chosen direction over a cardiac cycle.<br>Derive dimensional and functional parameters. | Left Ventricle (LV) chamber diameter at end-systole (LV ESD) and end-diastole (LV EDD) – measured in both long and short axis views.<br>Left Ventricle (LV) anterior and posterior wall thickness at end-systole (LV AWes, PWes) and end-diastole (LV AWed, PWed) – measured in both long and short axis view.<br>Heart Rate (HR). |
| Doppler Colour Flow Mapping | Visualize blood flow and its direction in different colors.<br>Qualitative demonstration of velocity distribution across heart chamber, orifice or vascular lumen.<br>Proper settings for the pulse repetition frequency (PRF)/velocity scale, gain and threshold of wall filtering. | Guidance for pulse wave Doppler velocity measurement at the sites of interest.<br>Visualize the abnormal flow patterns such high velocity flow jets. |
| Pulse Wave Doppler | Recording of blood flow velocity spectrum at site of interest.<br>Adjustable sample volume size to target vessel size.<br>Angle correction required for accurate measurement. Angle between ultrasound beam and flow less than 60 degrees. | Peak E and A velocity ratio of mitral inflow for left ventricular diastolic function.<br>Velocity Time Integral (VTI) of flow through the vascular lumen such as the ascending aorta or aortic orifice for calculating the flow rate.<br>Heart Rate (HR). |

All measures were obtained at least twice in different views/axes/traces (for example, from the left and right parasternal imaging windows or the short and long axis views of the heart). These measures were treated as duplicates/repeats in the data analysis.

Table 3 lists the cardiac parameters derived from the preliminary ultrasound measurements, and the formula for the related calculations.

**Table 3:**
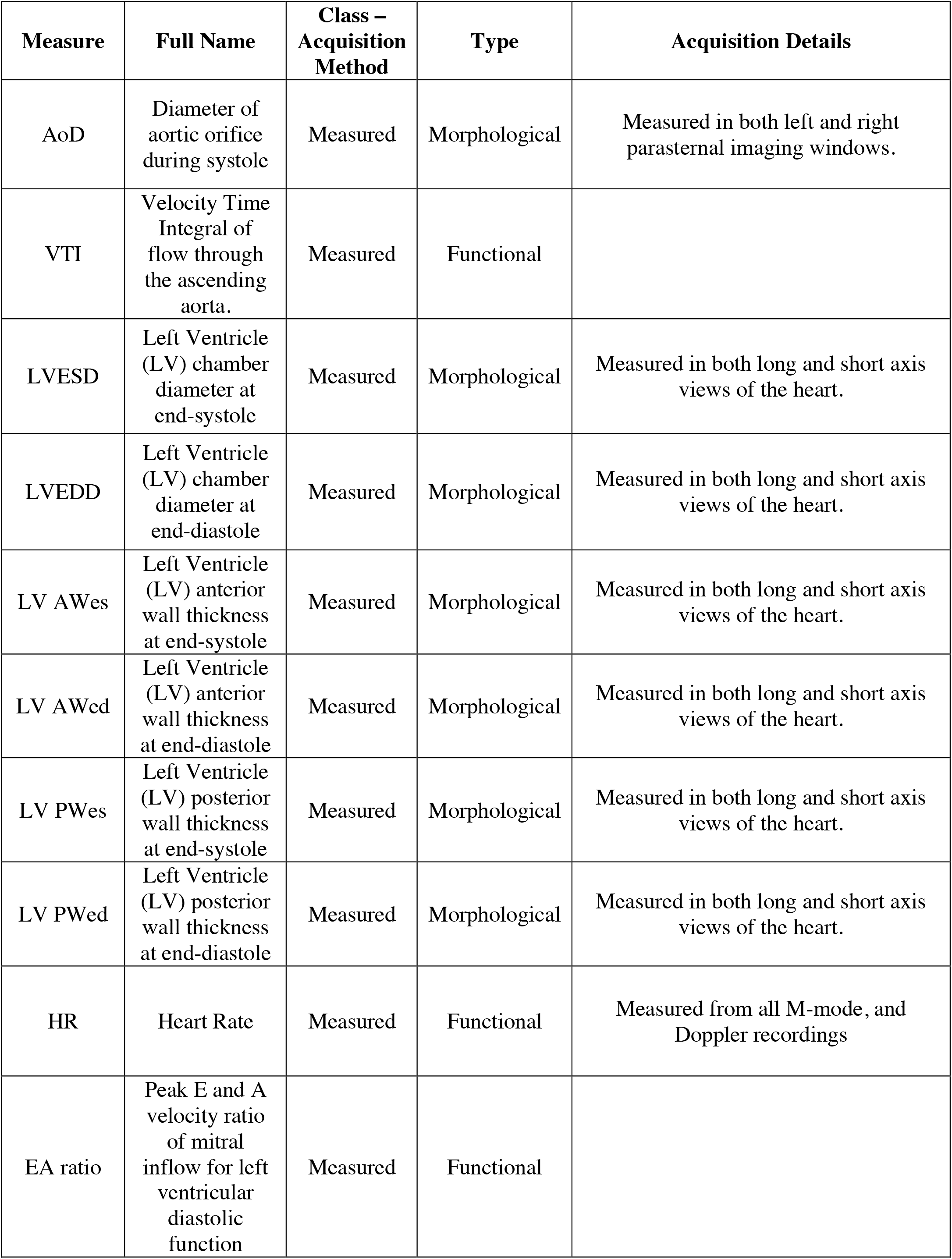
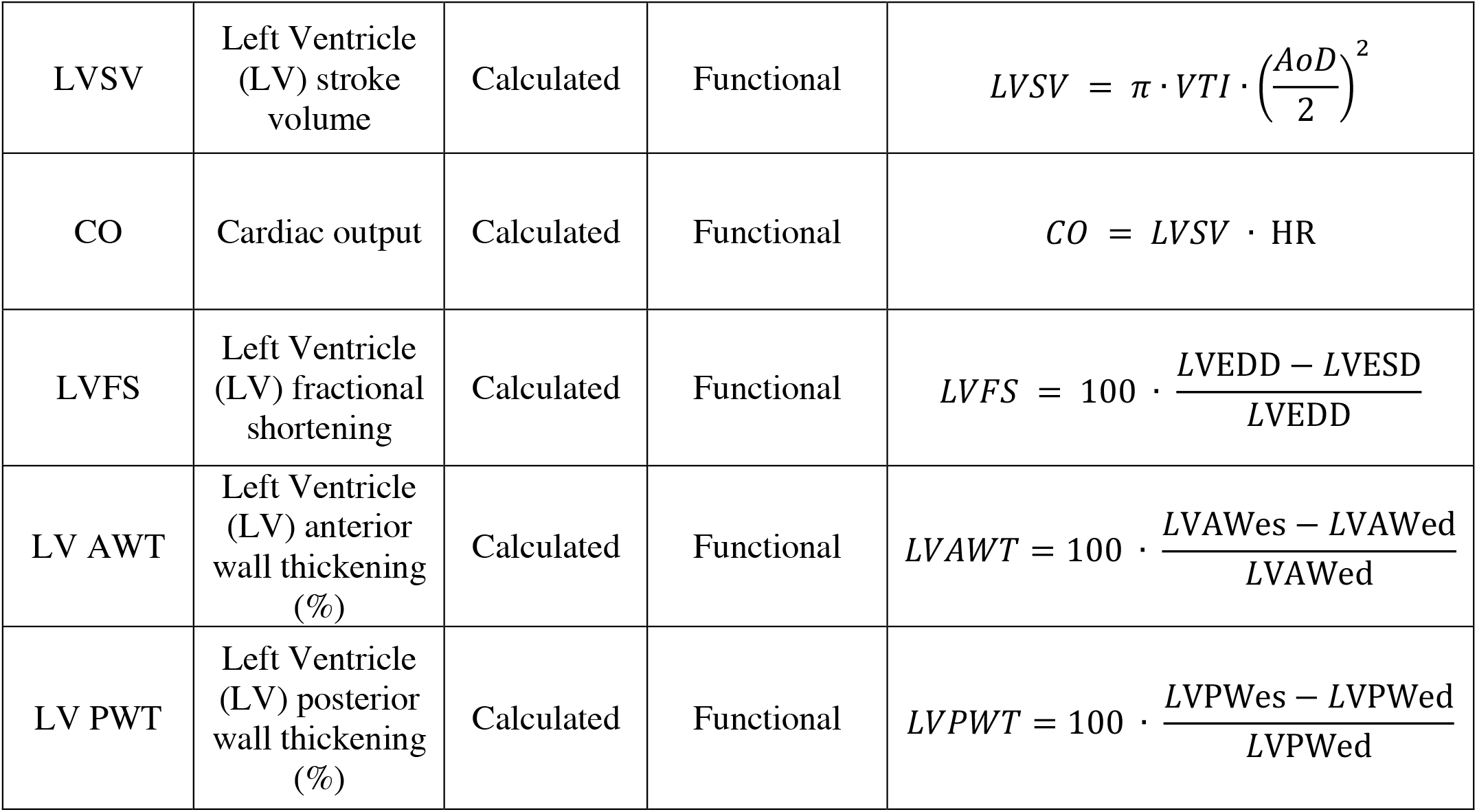
All the morphological and functional cardiac measures used in the study. The acronym, full name, and type (morphological or functional) of each measurement are given. For each measurement, the method of acquisition (termed “class”) (measured or calculated) is given along with the acquisition details (how it was measured (with ultrasound imaging) or calculated).

| Measure | Full Name | Class – Acquisition Method | Type | Acquisition Details |
| --- | --- | --- | --- | --- |
| AoD | Diameter of aortic orifice during systole | Measured | Morphological | Measured in both left and right parasternal imaging windows. |
| VTI | Velocity Time Integral of flow through the ascending aorta. | Measured | Functional |  |
| LVESD | Left Ventricle (LV) chamber diameter at end-systole | Measured | Morphological | Measured in both long and short axis views of the heart. |
| LVEDD | Left Ventricle (LV) chamber diameter at end-diastole | Measured | Morphological | Measured in both long and short axis views of the heart. |
| LV AWes | Left Ventricle (LV) anterior wall thickness at end-systole | Measured | Morphological | Measured in both long and short axis views of the heart. |
| LV AWed | Left Ventricle (LV) anterior wall thickness at end-diastole | Measured | Morphological | Measured in both long and short axis views of the heart. |
| LV PWes | Left Ventricle (LV) posterior wall thickness at end-systole | Measured | Morphological | Measured in both long and short axis views of the heart. |
| LV PWed | Left Ventricle (LV) posterior wall thickness at end-diastole | Measured | Morphological | Measured in both long and short axis views of the heart. |
| HR | Heart Rate | Measured | Functional | Measured from all M-mode, and Doppler recordings |
| EA ratio | Peak E and A velocity ratio of mitral inflow for left ventricular diastolic function | Measured | Functional |  |
| LVSV | Left Ventricle (LV) stroke volume | Calculated | Functional | $LVSV = \pi \cdot VTI \cdot \left(\frac{AoD}{2}\right)^2$ |
| CO | Cardiac output | Calculated | Functional | $CO = LVSV \cdot HR$ |
| LVFS | Left Ventricle (LV) fractional shortening | Calculated | Functional | $LVFS = 100 \cdot \frac{LVEDD - LVESD}{LVEDD}$ |
| LV AWT | Left Ventricle (LV) anterior wall thickening (%) | Calculated | Functional | $LVAWT = 100 \cdot \frac{LVAWes - LVAWed}{LVAWed}$ |
| LV PWT | Left Ventricle (LV) posterior wall thickening (%) | Calculated | Functional | $LVPWT = 100 \cdot \frac{LVPWes - LVPWed}{LVPWed}$ |

### Data analysis

Statistical analysis was performed in `R3.6.1` (48) using the `RStudio` interface (49) and the `brms` (50,51), loo (52) and `bayesplot` packages (53–56).

For each measure, we fit a Bayesian model with partial pooling to the *Z*-score values of our data, with mouse group and intercept as predictors and a random effect of mouse ID to account for redundancy in certain measures (measured more than once in different axes). We evaluated the need to account for a heteroscedasticity by comparing a model where variance of the posterior distribution varies between mouse groups (equation 1-A) versus a model with uniform variance across all groups (equation 1-B). All equations, including the priors used, can be found in equation 1.

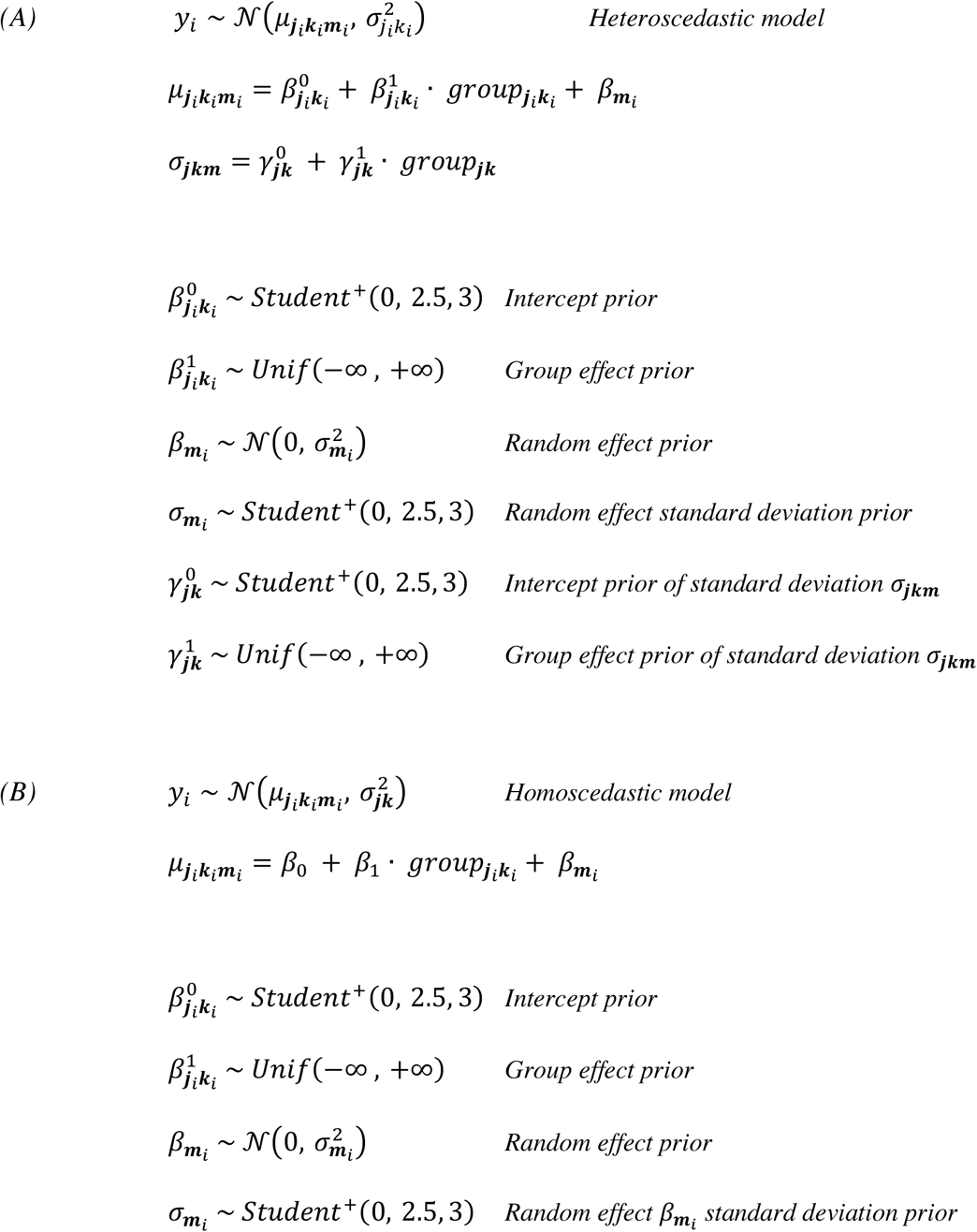

*Equation 1: (A) Heteroscedastic Bayesian model. (B) Homoscedastic Bayesian model. The Z-scored data for each observation (i) of every cardiac measure (j), mouse group (k) and mouse ID (m), y*_*i*_, *is modeled as a normal distribution with a mean* 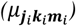 *and a variance which either varies with measure and mouse group* 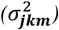 *(heteroscedastic) or varies only with measure* 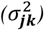 *(homoscedastic).* 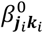 *is the intercept and* 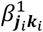 *is the group effect of the mean* 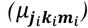. 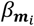 *is the random effect of mouse ID to account for repeated measurements and* 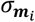 *is the standard deviation of the normal prior on* 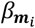. 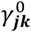 *is the intercept and* 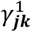 *is the group effect of the standard deviation* ***σ***_***jkm***_ *of y*_*i*_.*Student* ^+^(0, 2.5, 3) *is the half-student-t distribution with zero mean, scaling factor 2*.*5 and 3 degrees of freedom*.

The comparison of the two models was performed using the pareto-smoothed-importance-sampling leave-one-out cross-validation (*PSIS LOO CS*) method (57) in the loo package (52). For each measure, we then used the model that performed the best (Table 4).

**Table 4:** Type of Bayesian model used for each cardiac measure. One Bayesian model was run per cardiac measure comparing all genetic groups. For each model, the variance was either considered constant across genetic groups (homoscedastic) or varying between genetic groups (heteroscedastic).

| Cardiac Measure | Type of Bayesian Model |
| --- | --- |
| Aorta Diameter (AoD) | Heteroscedastic |
| Velocity Time Integral (VTI) | Homoscedastic |
| Peak E to A Velocity Ratio | Homoscedastic |
| Left Ventricular (LV) chamber diameter (end-systole) | Heteroscedastic |
| Left Ventricular (LV) chamber diameter (end-diastole) | Homoscedastic |
| Left Ventricular (LV) anterior wall thickness (end-systole) | Heteroscedastic |
| Left Ventricular (LV) anterior wall thickness (end-diastole) | Homoscedastic |
| Left Ventricular (LV) posterior wall thickness (end-systole) | Heteroscedastic |
| Left Ventricular (LV) posterior wall thickness (end-diastole) | Heteroscedastic |
| Left Ventricular (LV) anterior wall thickening | Homoscedastic |
| Left Ventricular (LV) posterior wall thickening | Heteroscedastic |
| Heart Rate (HR) | Heteroscedastic |
| Cardiac Output (CO) | Heteroscedastic |
| Left Ventricular Fractional Shortening (LVFS) | Homoscedastic |
| Left Ventricular Stroke Volume (LVSV) | Heteroscedastic |

For all sets of measures, we used the posterior distribution to obtain estimates for the mean for each genetic group. However, for measures fit using the heteroscedastic model, we obtained variance (sigma) for each genetic group, while measures fit using the homoscedastic model, we obtained a single posterior distribution for the variance (sigma) shared amongst genetic groups.

Once the posterior distributions were obtained a series of comparisons were performed. Specifically, each mutant group (total of 9) was compared to the WT control group for each cardiac measure (total of 15), for a total of 135 comparisons. Additionally, each mutant group (total of 9) was compared to every other mutant group per cardiac measure (total of 15), for a total of 540 comparisons.

For each posterior distribution we obtained the following metrics: median, mean, 89% credible intervals (CIs) and probability of direction (*pd*) (58,59). We controlled for the false discovery rate based on methods presented by J. Storey (60). Firstly, we calculated the posterior error probability (*PEP*) using: *PEP* = 1 − *pd*. PEP was used as a Bayesian p-value equivalent. Subsequently, we performed a correction on the PEP values using the formula:

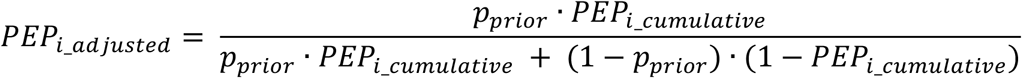

*Equation 2: Calculation of adjusted PEP of rank i to correct for multiple comparisons. p*_*prior*_ *is the prior estimate of the PEP value (p-value). Each PEP*_*i*_*cumulative*_ *is obtained after ranking the PEP values in increasing order (rank given by i) and adding all PEP values of less or equal rank to get PEP*_*I*_*cumulative*_.

The prior estimate of the *PEP* value (*p*_*prior*_) was obtained from the histogram of the *PEP* values (chosen bin size 0.05), using the following equation:

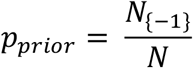

*Equation 3: Estimate of prior PEP value. N*_{−1}_ *is the sum of counts in all bins except for the first bin of the PEP histogram. N is the sum of counts in all bins. Chosen bin size was 0*.*05*.

*PEP*_*adjusted*_ ≤ 0.05 was considered to be of high confidence or significant.

### Principal Component Analysis and Total Correlation

Principal Component Analysis (PCA) and total correlation were used to explore the redundancy in our cardiac measures and genetic groups. This corresponds to comparing the genetic group by cardiac measure matrix of posterior means against its transpose. Firstly, for visual inspection, for each data matrix orientation (original and transposed), bootstrapping with replacement was performed to determine the median eigenvalue density for each PC component (as captured by the scree plot) along with the associated 95% confidence intervals. Secondly, to quantify the redundancy, total correlation was calculated (61).

Total correlation is sensitive to dimension (61). Thus, to make the total correlation comparable across both datasets, each dataset was iteratively downsampled to the lowest dimension (*N*_*genetic groups*_ × *N*_*genetic groups*_) by performing random selection without replacement of columns or rows where appropriate. For each dataset, the mean and standard deviation of the total correlation across all iterations were calculated. Further details on the total correlation calculation can be found in the Supplementary Materials.

## Results

The type of model (heteroscedastic or homoscedastic) run per cardiac measure is shown in Table 4. Depending on the model, coefficient estimates (posterior distributions) were obtained either only for the median or for both the median and variance of the coefficients. The coefficient estimates for the median are shown in Figure 1.

**Table 4:** Total correlation values per dataset. First column is the total correlation of the original datasets. In the second and third column are the mean total correlation and standard deviation respectively, across all iterations, for the downsampled data.

|  | Total Correlation<br>Original Dimensions | Total Correlation<br>Downsampled Data | Total Correlation<br>Downsampled Data<br>Standard Deviation |
| --- | --- | --- | --- |
| Original-Cardiac Measures | 66.18 | 18.15 | 1.3 |
| Original-Genetic Groups | 10.79 | 19.36 | 1.26 |

**Figure 1:**
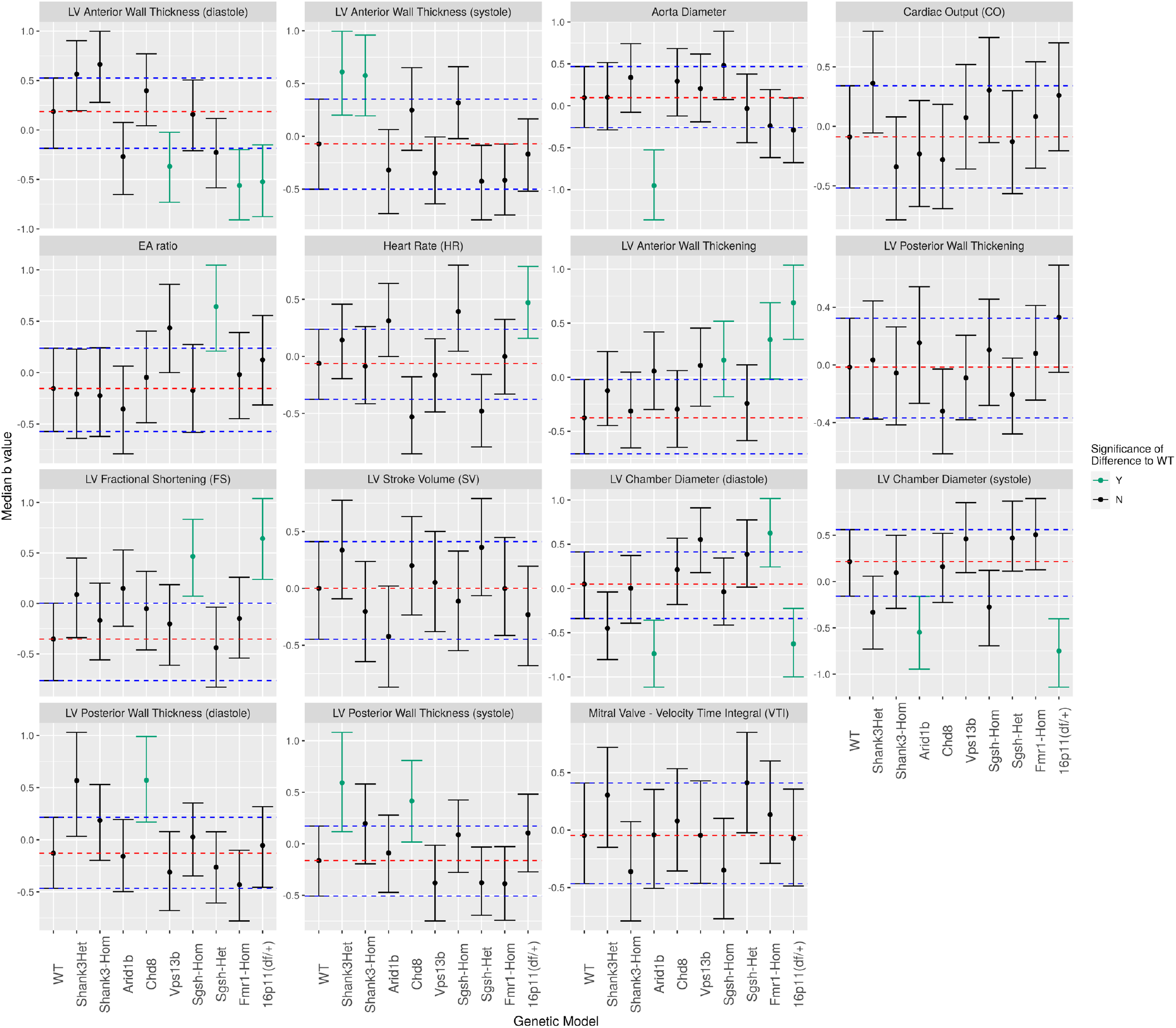
Median coefficient estimates (b-values) per genetic group per cardiac measure. Unadjusted 89% Credible Intervals (CIs) shown as whiskers. The blue dashed lines indicate the upper and lower 89% CIs for the WT control group. The red dashed line indicates the median of the WT control group. Green colour indicates the groups that are significantly different from WT controls. Significance assigned based on FDR corrected q values.

We determined the number of “high confidence” or “significant” (*PEP*_*adjusted*_ ≤ 0.05) tests per genetic group, compared to all tests performed for that group. The fraction of high-confidence tests was used as a metric of difference compared to the reference group. The higher the fraction, the more (measures with) significant differences between the groups. Similarly, for each cardiac measure the number of high-confidence tests was calculated (relative to the total number of tests). In this case, it was interpreted as the prevalence of each cardiac measure in an abnormal cardiac phenotype. The higher the fraction, the more often would that cardiac measure be abnormal (relative to the reference group).

### Comparing ASD-related genetic models to littermate wild-type (WT) controls

From our results, the differences between mutant groups and WT controls varied between mutant groups in both number and magnitude. The mutant group with the greatest number of significant differences in cardiac phenotype relative to WT controls was the 16p11.2 (deletion) group (6/15 measures) with the *Arid1b* ^*(+/-)*^ and *Fmr1*^*(-/-)*^ groups following (3/15 measures). Next were the *Shank3*^*(+/-)*^ *(Δexon 4-9), Chd8*^*(+/-)*^ and *Sgsh*^*(-/-)*^ groups (2/15 measures each but not in all the same cardiac measures) (Figure 3A). In all comparisons to WT controls, the cardiac measures that were most often found abnormal were, the left ventricular (LV) anterior wall thickening (LVAWT) (3/9 groups), LV chamber diameter at end-diastole (LVEDD) (3/9 groups), and LV anterior wall thickness at end-diastole (LVAWed) (3/9 groups) and end-systole (LVAWes) (2/9 groups) (Figure 3B). Overall, there were mainly morphological differences, but functional differences were also present.

**Figure 2:**
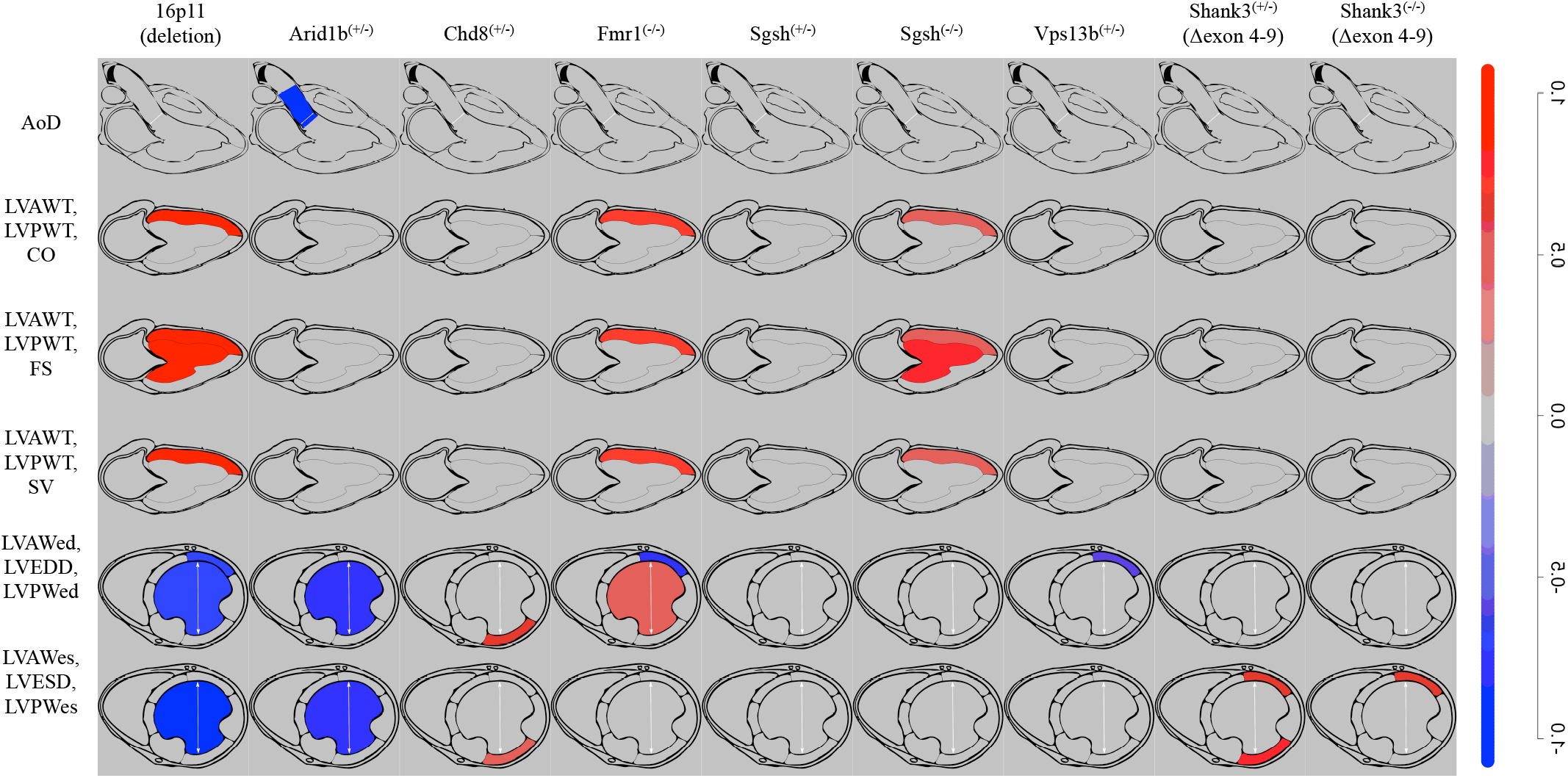
Normalised median differences per cardiac measure between each mutant mouse group and wild-type controls (WTs). The colour gradient was generated mapping to the range of normalised median values; only the data from the high-confidence (significant) tests across all cardiac measures and groups were used. For all other (non-significant) tests, a grey colour was assigned, same as the background. Each column corresponds to a mutant mouse group and each row depicts one or a set of cardiac measures. Abbreviations: AoD – Aorta Diameter; LVAWT – left ventricular anterior wall thickening; LVPWT – left ventricular posterior wall thickening; CO – cardiac output; FS – fractional shortening; LVAWed – left ventricular end-diastolic anterior wall thickness; LVEDD – left ventricular end-diastolic (chamber) diameter; LVPW – left ventricular end-diastolic posterior wall thickness; LVAWes – left ventricular end-systolic anterior wall thickness; LVESD – left ventricular end-systolic (chamber) diameter; LVPWes – left ventricular end-systolic posterior wall thickness.

**Figure 3:**
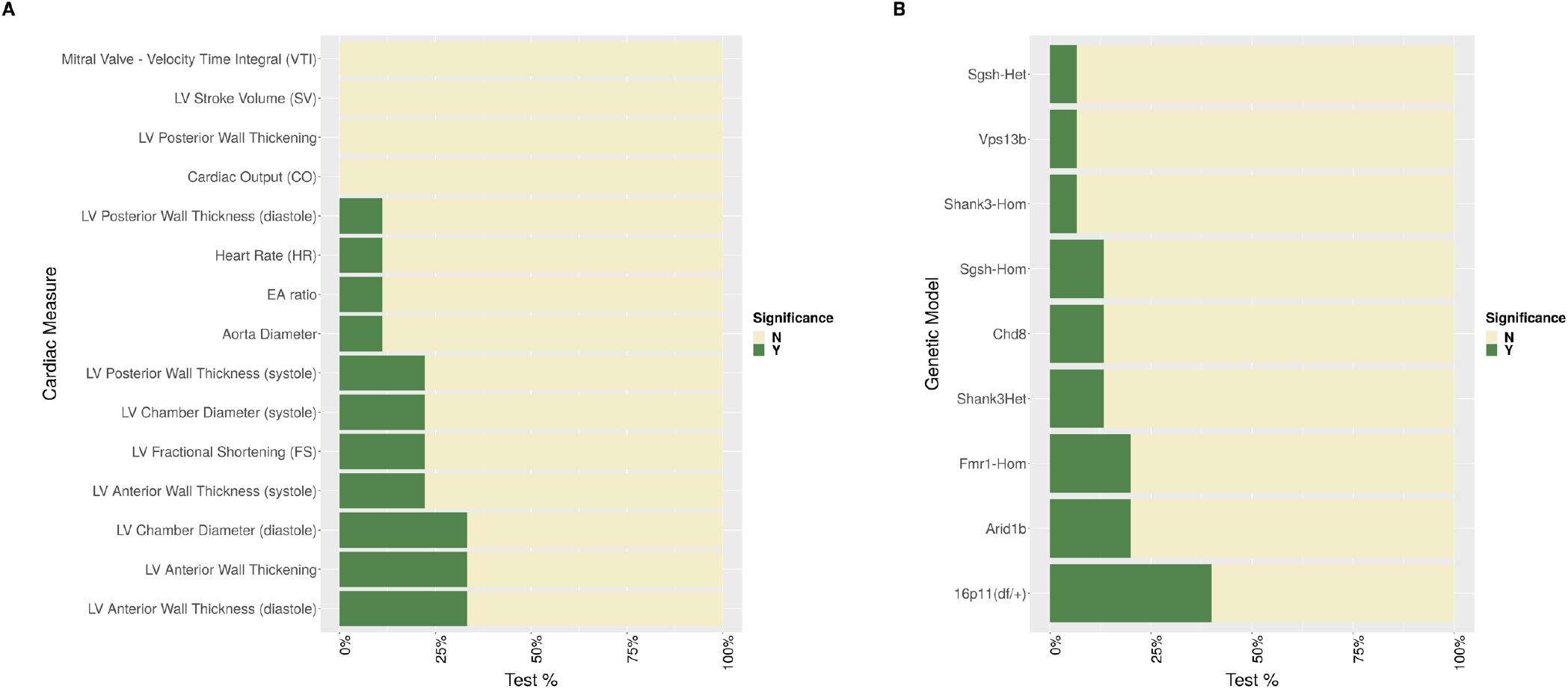
Barplots indicating the percent of significant (high confidence) comparisons (tests) between mutants and WT controls. (A) For each cardiac measure all ASD-related mutant groups were compared to WT controls; the same total number of comparisons (tests) was conducted for all cardiac measures. The green bar indicates the percent of significant comparisons (tests) for each cardiac measure. (B) Similar to (A), each ASD-related mutant group was compared to WT controls for all cardiac measures; same number of comparisons (tests) was conducted for all mutant groups. The green bar indicates the percent of significant comparisons (tests) for each group.

### Comparing ASD-related genetic mouse groups

To explore the heterogeneity amongst mutant groups, we identified the cardiac measures that differed most between groups (Figure 4A), the mutant groups with most differences from all other groups (Figure 4B), and the mutant group pairs with the most differences.

**Figure 4:**
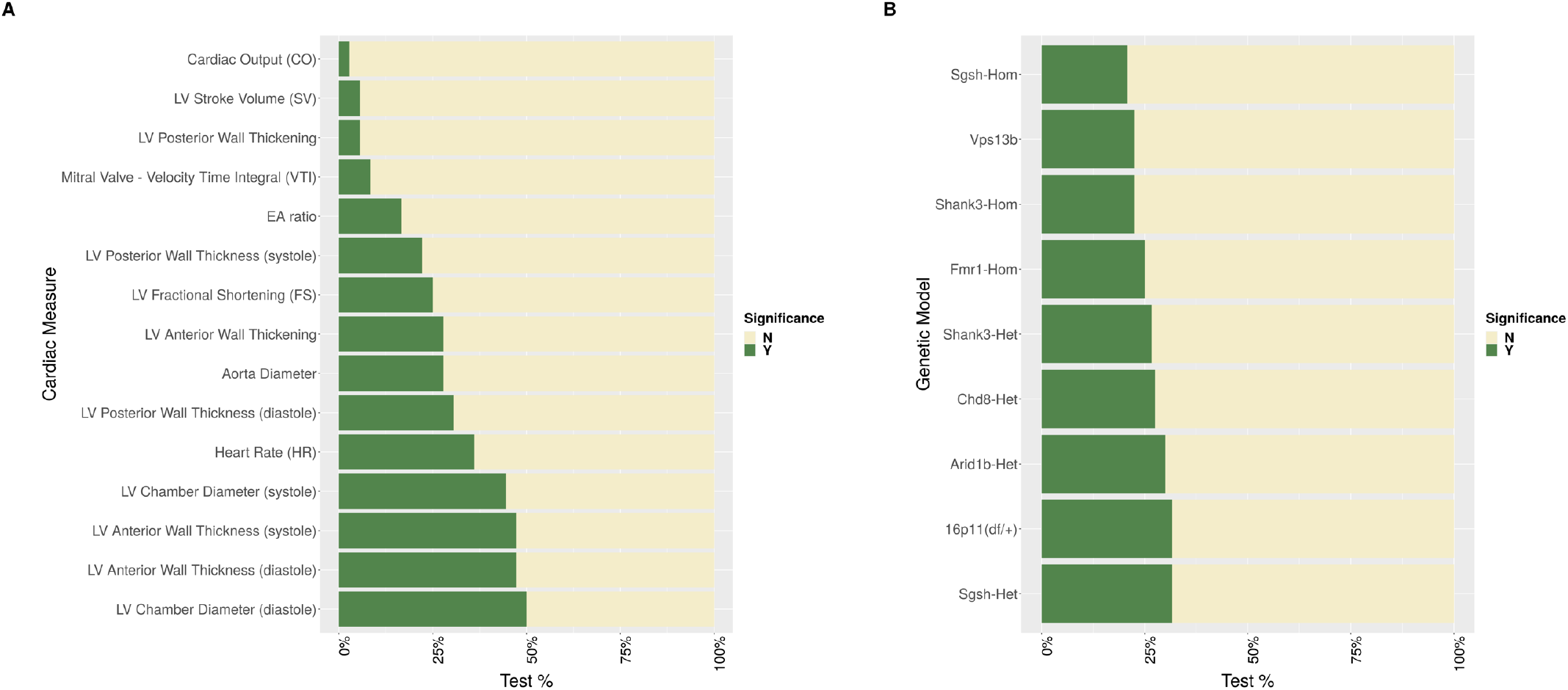
Barplots indicating the percent of significant (high confidence) comparisons (tests) between all ASD-related genetic mutant groups. (A) For each cardiac measure all ASD-related mutant groups were compared to each other; the same total number of comparisons (tests) was conducted for all cardiac measures. The green bar indicates the percent of significant comparisons (tests) for each cardiac measure. (B) Similar to (A), each ASD-related mutant group was compared to every other mutant group, for all cardiac measures; same number of comparisons (tests) was conducted for all mutant groups. The green bar indicates the percent of significant comparisons (tests) for each group.

A greater number of significant differences was present between mutant groups than when mutant groups were compared to WT controls. The measures driving the inter-group variation tended to be associated more with morphological changes than functional changes (Figure 4A). Specifically, the mutant groups firstly differ in LV chamber diameter at end-diastole (LVEDD) (significance found in 18/138 pairwise comparisons) and, secondly, in LV anterior wall thickness at end-systole and end-diastole (LVAWes, LVAWed) (17/138 pairwise comparisons each). Thirdly they differ in LV chamber diameter at end-systole (LVESD) (16/138 pairwise comparisons), followed by heart rate (HR) (13/138 pairwise comparisons). The 16p11.2 (deletion) and *Sgsh*^*(+/-)*^ groups had the most differences from all other mutant groups (38/1080 pairwise comparisons), followed by the *Arid1b*^*(+/-)*^ (36/120 pairwise comparisons) and *Chd8*^*(+/-)*^ (33/120 pairwise comparisons) groups (Figure 4B).

Conducting pairwise mutant group analysis, the *Shank3*^*(+/-)*^ *(Δexon 4-9)* - *Sgsh*^*(+/-)*^ and the *Chd8*^*(+/-)*^ - 16p11.2 (deletion) pairs had the greatest number of significant differences in cardiac measures (8/15 measures), followed by the *Shank3*^*(-/-)*^ *(Δexon 4-9)* - 16p11.2 (deletion) and *Arid1b*^*(+/-)*^ - *Sgsh*^*(+/-)*^ pairs (7/15 measures each).

Of interest is the comparison of the distributions in Figures 3A and 4A. When compared to each other, mutant groups tended to differ more in morphological measures than in functional measures (Figure 4A), but when compared to WT controls, they tended to differ roughly equally in both (Figure 3A).

### Principal Component Analysis and Total Correlation

For the 15 cardiac measures, PCA revealed that 9 principal components were required to account for ∼99% of the variance in the data. This can be explained by the fact that, from the set of 15 measures, 5 were not directly measured but were calculated using others, as described in Methods. For the 10 genetic groups, PCA also revealed that 9 principal components were required to account for ∼99% of the variance in the data. Therefore, there doesn’t seem to be any pattern of similarity (redundancy) between the mutant groups with regard to cardiac phenotype.

**Figure 2:**
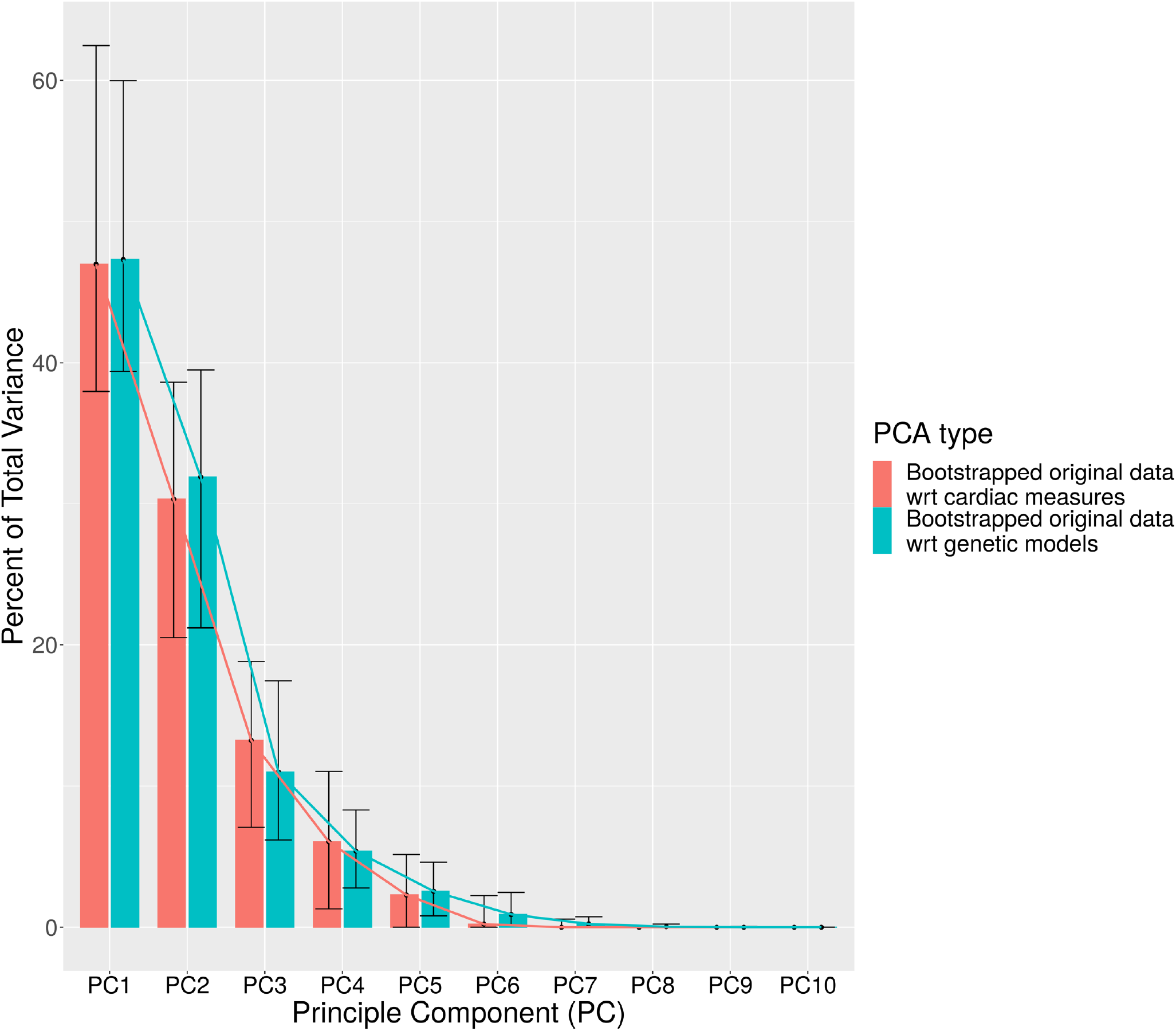
Scree plot of PCA on cardiac measures and PCA on genetic models. Bootstrapping with replacement of the data was performed to determine the median eigenvalue density for each PC component along with the associated 95% confidence intervals. The bootstrapped data are shown along with the bootstrapped median values and 95% confidence intervals. The estimated decay curve is also mapped.

To inspect the comparable redundancy between cardiac measures and genetic groups, the total correlation was computed (Table 4). The total correlation of 15 cardiac measures was 66.18 and of 10 genetic groups was 10.79. As total correlation is sensitive to the number of dimensions, we randomly downsampled both datasets to 10 dimensions with 1000 permutations (reporting mean +/- standard deviation). The total correlation of the downsampled cardiac measures was 18.15 +/- 1.3 and of the downsampled genetic groups was 19.36 +/- 1.26. Thus, the total correlation, and therefore redundancy, is comparable between cardiac measures and genetic groups. Amongst cardiac measures and amongst genetic groups, respectively, some redundancy or correlation exists since for random data the total correlation would be zero.

## Discussion

Our analysis revealed small-scale alterations in cardiac structure and function in ASD compared to WT, mirroring the clinical reports. However, significant differences of note were present. Firstly, the alterations each mutant group presented to WT controls were largely not consistent between groups. Therefore, the heterogeneity characterising ASD in other phenotypes (7,8) is recapitulated here. Specifically, when compared to each other, mutant groups presented a greater number of significant differences than when compared to WT controls. As mentioned, this is possible due to certain mutant groups differing insignificantly from WT controls in an opposite fashion (for the same cardiac measure). Secondly, mutant groups tended to differ primarily in measures of LV structure, while when compared to WT controls, they tended differ in both morphological and functional measures, with a small prevalence of the former. However, there was overlap in the overall associated measures with HR and LV chamber diameter and anterior wall thickness being strong contributors in both cases. Moving forward, it would be of interest to expand this work to include more ASD-related genetic models and further assess the emerging patterns.

As previously mentioned, the available clinical literature on the cardiac phenotype of ASD patients is not extensive and not always consistent. Nevertheless, with our standardised protocol we interrogated the cardiac phenotype of our genetic models and compared our findings to the clinically documented abnormalities associated with the corresponding ASD genetic mutations. This information can be found in the Supplementary Materials. Overall, our screening process captures the abnormalities reported clinically for each genetic mutation. For example, in the *Arid1b* ^*(+/-)*^ group, in agreement with the clinical observations, we observed arterial stenosis at the level of the aortic valve. We also see evidence of the common cardiac abnormalities associated with ASD, for example autonomic dysregulation (manifested as abnormal HR). A detailed per-model account of this comparison can be found in the Supplementary Materials.

Given the heterogeneity observed, it is valuable to determine how it compares to a standardised categorisation of our genetic groups. One such categorisation can be found in the SFARI Gene database (Simons Foundation; http://gene.sfari.org/) (62) which has assigned a score to each genetic variant quantifying how closely each genetic group is linked to ASD. According to the SFARI Gene database, the 16p11.2 (deletion), *Arid1b*^(+/-)^, *Fmr1*^(-/-)^, *Shank3 (Δexon 4-9), Chd8*^(+/-)^, *Vps13b*^(+/-)^ groups hold a high score and the *Sgsh* group holds a lower score. However, even when assessing our results with the SFARI score in mind, a pattern cannot be identified. As shown in Figure 3B, considering the number of differences to WT controls, certain groups with a high SFARI score rank high (16p11.2 (deletion), *Arid1b*^(+/-)^, *Fmr1*^(-/-)^, *Shank3 (Δexon 4-9)*), while others rank low (*Chd8*^(+/-)^, *Vps13b*^(+/-)^). The *Sgsh* group with a low SFARI score ranks somewhere between the two. This discrepancy between SFARI gene score and severity of cardiac phenotype in each group could be driven by different expression levels of the associated gene between the brain and the heart.

Consequently, it seems unfeasible, with this type of assessment, to identify an overarching ASD-related cardiac phenotype, shared between all individual genetic mutant groups and setting them apart from WT controls. It seems more likely that there would be subgroups of ASD-related models, potentially not based on how closely they are linked to ASD (SFARI Gene score), each with a different cardiac phenotype compared to WT controls. This is something to be explored further in the future.

Knowing the abnormalities present, allows us to choose alternate measures to explore more closely the observed changes and probe the underlying mechanisms. Additionally, categorising measures as morphological or functional can offer insight about common etiology of the models’ cardiac phenotype. For example, the Sinoatrial Node (SAN) is the principal pacemaker of the heart, it is characterised by cellular heterogeneity and its formation and function are controlled by a series of upstream factors (63,64). Its cells have been clustered based on their downstream function, revealing a wide range of targets (64). We could then hypothesise that a disruption affecting the formation of SAN (starting at embryonic day 9.5) could result in a spectrum of functional abnormalities across various genetic groups (depending on the origin and type of the disruption). Similarly for structural abnormalities.

The cardiac phenotype of each of our ASD mouse mutant groups was consistent with results seen in the corresponding human population to the extent that a direct comparison is possible (see supplementary materials). For example, in the *Arid1b* ^*(+/-)*^ group, in agreement with the corresponding clinical reports (5), we observed arterial stenosis at the level of the aortic valve. Similarly, the cardiac abnormalities associated with ASD as a whole (as reported in literature) are also observed across all our ASD groups. For example, autonomic regulation is impaired in the autism population which is consistent with our observations that HR was one of the main differing measures between ASD mutant groups and WT controls.

Finally, Principal Component Analysis (PCA) and total correlation comparison revealed a comparable redundancy between the cardiac measures and the genetic groups, while ensuring the outcome wasn’t solely driven by underlying noise. One would expect cardiac measures to have a larger redundancy (since all measures are related to the heart), compared to the genetic groups (each was created to carry a different genetic modification). However, this result indicates that the cardiac measures used in this study are more independent than one would expect (with some correlation amongst them still present). Thus, once again, confirming that our UBM protocol captures various dimensions of cardiac functionality, as intended, and can be used as an effective screening protocol. For the genetic groups, this result indicates there is some correlation between the ASD-related models but overall, as mentioned previously, the heterogeneity characteristic of ASD is recapitulated in the cardiac phenotype as well.

## Limitations

There are certain limitations to this work which must be considered. Firstly, this work was conducted using adult mice. However, various genetically driven cardiac abnormalities, and especially CHD, occur early in life. So, by screening in adult age, there is a risk of observing a modified state which resulted from various adaptive mechanisms acting throughout development. Secondly, only male mice were used in the study. Given the known sex differences in ASD (63) and cardiac function (64), we recognise it may be of value to repeat this work on female mice. Thirdly, the downsampling procedure used for the calculation of total correlation is not a standard approach for comparing two datasets of differing dimensions. So further investigation is needed to verify this approach. Finally, a direct comparison between our results and the clinical literature is not possible because clinical studies often employ different study designs (assessment protocols, sample sizes etc) and recruit both male and female patients. Namely, in this study we did not perform direct assessment of the atrioventricular septum, the right side of the heart, the extracardiac space or the conduction system of the heart, which are widely mentioned in many of the clinical reports, as is shown in Table 1 of the Supplementary Materials. However, potential links can be inferred based on knowledge of the overall structure and function of the cardiovascular system. As the number of ASD-related genetic models screened with this protocol increases and as the corresponding clinical data amounts, we will be able to make even more concrete comparisons.

## Conclusion

This work sheds light on the spectrum of cardiac abnormalities associated with ASD-related genetic abnormalities. The heterogeneity characterising ASD in other phenotypes is recapitulated here, with more differences seen between mutant groups than when compared to WTs. Alterations were small in scale, but significant, and further exploration of more models is needed to validate the observed patterns. Our high frequency ultrasound imaging protocol as a method of cardiac assessment provides a well-rounded view of cardiac structure and function, effectively capturing the clinically reported cardiac abnormalities. Thus, it can be used as a screening protocol moving forward. The detected cardiac abnormalities can then be further examined using potentially more sensitive methods to explore their underlying mechanisms. By classifying cardiac measures as morphological or functional, the etiology of mutant phenotype can be better understood, and any common underlying mechanisms can be elucidated. Clinically, knowledge of the cardiac abnormalities associated with ASD can be greatly beneficial as, even non-lethal cardiac abnormalities can impact the normal development and function of various other biological systems, such as the brain. In addition, the presence of specific cardiac abnormalities may provide mechanistic insights for a patient’s ASD subtype.

## Abbreviations

ASD: Autism Spectrum Disorder
CHD: Congenital Heart Disease
UBM: Ultrasound Bio-Microscopy
LV: Left Ventricle
WT: Wild-Type
HET: Heterozygous (+/-)
HOM: Homozygous (-/-)
AoD: Aorta Diameter
HR: Heart Rate
LVAWed: Left Ventricular Anterior Wall Thickness at End-Diastole
LVAWes: Left Ventricular Anterior Wall Thickness at End-Systole
LVEDD: Left Ventricular Chamber Diameter at End-Diastole
LVESD: Left Ventricular Chamber Diameter at End-Systole
LVPWed: Left Ventricular Posterior Wall Thickness at End-Diastole
LVPWes: Left Ventricular Posterior Wall Thickness at End-Systole
LVAWT: Left Ventricular Anterior Wall Thickening
LVPWT: Left Ventricular Posterior Wall Thickening
FS: Fractional Shortening
CO: Cardiac Output
LVSV: Left Ventricular Stroke Volume
sys: systole
di: diastole
PCA: Principal Component Analysis
MRI: Magnetic Resonance Imaging
ECG: Electrocardiography

