## Supplementary Materials for "Genetic Mouse Models of Autism Spectrum Disorder Present Subtle Heterogenous Cardiac Abnormalities"

In depth comparison of Results to Clinical Reports

As previously mentioned, the available clinical literature on the cardiac phenotype of ASD patients is sparse and not always consistent. Nevertheless, in Table 1 we collected the most frequently documented cardiac phenotypes associated with the ASD genetic mutations corresponding to the genetic models of our study. Thus, we were able to assess whether our screening process captures any documented abnormalities. It must be noted that an association between the documented clinical data and the results of this study cannot be direct, primarily because different measures were used. Namely, in this study we did not perform direct assessment of the atrioventricular septum, the right side of the heart, the extracardiac space or the conduction system of the heart, which are widely mentioned in many of the clinical reports. However, hypotheses on the potential links can be made based on the overall structure and function of the cardiovascular system. Elaborating on the data shown in Table 1, the potential association between our results and the clinically observed cardiac phenotype for each genetic mutation can be seen below:

*Shank3 (Δexon 4-9)*

In this study the *Shank3*$^{(+/-)}$ *(Δexon 4-9)* and *Shank3*$^{(-/-)}$ *(Δexon 4-9)* mutant groups were used. For each group we observed significant differences in the LV structure compared to WT controls. Specifically, the *Shank3*$^{(+/-)}$ *(Δexon 4-9)* group showed an increased LV posterior wall thickness at end-diastole (LVPWed), while both groups showed an increased LV anterior wall thickness at end-systole (LVAWes).

Molecular analysis has shown abundant expression of Shank3 mRNA in the heart (65,66). Clinically, the most consistently documented cardiac abnormalities associated with the Shank3 microdeletion (Phelan McDermid syndrome) are Patent Ductus Arteriosus (PDA), tricuspid valve regurgitation, Atrial Septal Defect (ASD) and Total Anomalous Pulmonary Venus Return (TAPVR) (5,66). The prevalence of these abnormal cardiac phenotypes in patients is larger than 25% (67). In mice, Shank3 has been found to play a role in myocardial remodeling following myocardial infarction (68).

The presence of a patent ductus arteriosus could lead to pulmonary hypertension which, causing an increased load on the ventricles, can in turn lead to an enlarged LVAWes. Similar findings can be caused from a left atrial septal defect or a dysfunction of the right side of the heart (for example in tricuspid valve regurgitation) which also increase the load on the LV (69–71).

In the clinical literature there was no discrimination between heterozygous (+/-) and homozygous (-/-) patients carrying the Shank3 mutation, so all the clinical information was taken as a reference point for both groups.

*Fmr1*$^{(-/-)}$

In the *Fmr1*$^{(-/-)}$ group we observed an increased LV chamber diameter at end-diastole (LVEDD) and decreased LV anterior wall thickness at end-diastole (LVAWed), as well as an increased LV anterior wall thickening (LVAWT), compared to WT controls. One of the main phenotypes observed clinically in patients carrying the Fmr1 mutation is mitral valve prolapse (MVP) (5). A dysfunction of the mitral valve connecting the left atrium (LA) to the LV, such as MVP, can result in excess load on the left ventricle and in turn result in increased chamber diameter. In mild cases of MVP, LV attains a compensated normal function by an increase in LVEDD while LVESD (LV end-systole diameter) remains the same (72–74). Additionally, autonomic dysfunction has been documented (75) which could be expressed as a change in LV anterior wall thickening (LVAWT). According to clinical literature, the prevalence of congenital heart disease in Fmr1 is less than 10% (5).

*Arid1b^(+/-)*

The ARID1B gene, besides being a prominent autism-related gene, is also associated with Congenital Heart Disease (CHD) (4) with a prevalence in 20-40% of CHD cases (5).

In the *Arid1b^(+/-)* group, in agreement with the clinical observations, we observed arterial stenosis at the level of the aortic valve. We also observed a decreased LV chamber diameter both an end-systole and end-diastole (LVESD, LVEDD). This observation can be linked to certain clinically reported phenotypes in patients with an Arid1b mutation (Table 1). Specifically, pulmonary stenosis (5) inevitably results in an increased right ventricular (RV) wall or chamber to compensate for the increased afterload. It is known that changes in the structure and pressure of the RV can lead to changes in the LV through pressure gradients and/or morphological changes in the intraventricular septum (IVS). For example, an increased RV chamber can cause a flattening of the IVS towards the LV, thus inevitably causing a decrease in LV chamber diameter (69,70).

As previously stated, a lack of assessment of the right side of the heart and the IVS, doesn't allow us to be more assertive of our interpretations.

*16p11.2 (deletion)*

In the 16p11.2 (deletion) group we observed decreased LV chamber diameter at end-systole and end-diastole (LVESD, LVEDD) and decreased LV anterior wall thickness at end-diastole (LVAWed). We also observed an increased LV anterior wall thickening (LVAWT) and fractional shortening (FS). Additionally, we observed an increased heart rate (HR).

Clinically, Tetralogy of Fallot (ToF) and associated anomalies have been observed (76). The primary abnormality of ToF is pulmonary stenosis. As previously mentioned, pulmonary stenosis increases the afterload for the RV causing increased stress and morphological changes. These RV changes can induce changes in the LV consistent with a decrease in chamber diameter (as was also seen in the Arid1b group). Similar effects could be caused by pulmonary valve regurgitation also clinically observed as an associated anomaly of ToF (69–71). The induced stress from the RV to the LV can also affect myocardial function through the cardiac cycle thus causing changes in LVAWT and FS (73).

Additionally, genes harbored in the 16p11.2 microdeletion have been linked to aortic valve development (77). Inevitably, abnormal aortic valve function could result in LV changes due to changes in afterload. Aortic valve dysfunction can result in increased afterload, leading to LV remodeling and increased HR. Both are consistent with our findings.

It must be noted that, contrary to traditional practices and views (78), FS (just like ejection fraction (EJ)) has recently been found to be insensitive to mild diastolic dysfunction as it is rather preload and afterload dependant (73). Even more, FS (and EJ) has been shown to overestimate systolic function in cases of concentric hypertrophy or LV remodeling making it a less reliable measure of systolic function (79). For example, should some LV remodeling have occurred due to atrioventricular septal defects or pressure gradients from the RV (all possible phenotypes reported clinically), then a change in FS could be reported even if a change in systolic function (signaling) hasn't occurred (69,80).

*Chd8*$^{(+/-)}$

In the *Chd8*$^{(+/-)}$ groups not many significant differences compared to WT controls were identified. Specifically, we only found increased LV posterior wall thickness at end-systole (LVPWes) and end-diastole (LVPWed).

The literature available on the cardiac phenotype of patients with a Chd8 mutation is limited more so than for the other mutations. However, information has accrued on the cellular and molecular level linking the CHD8 protein to cardiac development. Firstly, CHD8 plays a role in cardiac development through the Wnt signaling pathway, specifically, through the regulation of β-catenin (81). Studies in the mouse have shown β-catenin contributes to the regulation of postnatal hypertrophic growth by acting upstream of fibroblast growth factor (FGF) signaling. FGF signaling regulates second heart field (SHF) progenitors which give rise to the atrial, ventricular and outflow tract structures (82,83). Secondly, in the mouse, Chd8 mRNA has been detected in high levels during early embryogenesis and up to 10 days postnatally with a drop-off in concentration thereafter, while complete knockout of Chd8 leads to *in utero* embryonic demise following sever hemorrhage, indicative of cardiovascular defects (84). Thirdly, in the rat, Chd8 has been detected during embryonic and postnatal cardiac development in myocytes and fibroblasts. Even more, Chd8 has been proposed to be a new AKAP (A-kinase anchoring protein) in humans. Defects in the AKAP function have been shown to lead to arrythmias, hypertrophy and eventual progression to heart failure (85). Given the above, our findings of a reduced heart rate (HR) and LV remodeling (increased LVPWED) are supported.

Of note is the indirect link of CHD8 to cardiac dysfunction in CHARGE syndrome. CHARGE syndrome is characterised by a dysfunction of the CHD7 gene. However, CHD8 is an interacting partner of the CHD7 protein. Changes in interacting proteins necessary for the function of CHD7, such as CHD8, have been proposed as an underlying cause of the disease

(86). The main cardiac abnormalities in CHARGE syndrome are Tetralogy of Fallot (ToF), Patent Ductus Arteriosus, double outlet RV with AV canal, Ventricular Septal Defect, Atrial Septal Defect (w/ or w/o cleft mitral valve)(30). As mentioned for the 16p11.2 (deletion) group, ToF alone can result in HR abnormalities and LV remodeling.

*Sgsh*

As for the *Shank3 (Δexon 4-9)* model, for the *Sgsh* model we had a *Sgsh*$^{(+/-)}$ and a *Sgsh*$^{(-/-)}$ group, each of which was compared to WT controls. The *Sgsh*$^{(+/-)}$ group showed an increase in the E/A ratio. The *Sgsh*$^{(-/-)}$ group showed an increase in LV anterior wall thickening (LVAWT) and fractional shortening (FS). Overall, clinically, there is no significant signs of cardiac disease. However, the main cardiac abnormality is congestive cardiomyopathy which eventually results in enlarged LV. Other reported cardiac abnormalities include valvular abnormalities (mitral and aortic valves), LV systolic and diastolic dysfunction, from subclinical (with normal ejection fraction (EF)) to more severe (with reduced EF). There have also been reports of reduced E/A ratio (due to increase A wave amplitude) (87–89).

Our results for both groups align with the more common subclinical LV systolic/diastolic dysfunction since the EF is unchanged. Additionally, all changes involve measures directly linked to signaling during systole and diastole.

It is important to note once again, as for the 16p11.2 (deletion) group, that there is accumulating evidence that EF (just like FS) is not a reliable marker of systolic function in certain cases of LV remodeling and hypertrophy and other measures should be used (for example measures of myocardial strain) (79).

*Vps13b*$^{(+/-)}$

The *Vps13b*$^{(+/-)}$ group is the one with the fewest differences in cardiac phenotype compared to WT controls. Specifically, a decrease in LV anterior wall thickness at end-diastole (LVAWed) was observed. This closely mirrors the literature where clinically a normal heart anatomy (aorta diameter, septum and LV posterior wall thickness) is primarily reported. The abnormal phenotypes reported most consistently mainly are essential and pulmonary hypertension and cardiac systolic murmurs (90–92).

| Genetic Model | Associated Syndrome | Clinically Documented Phenotype | Observed Significant Phenotype | Association | Reference |
|---|---|---|---|---|---|
| ***Shank3*$^{(+/-)}$ *(Δexon 4-9)* *Shank3*$^{(-/-)}$ *(Δexon 4-9)*** | Phelan McDermid Syndrome | • Congenital Heart Disease prevalence >25%<br>• SHANK3 mRNA abundantly expressed in the heart<br>• Tricuspid Valve Regurgitation<br>• Atrial Septal Defect (ASD)<br>• Patent Ductus Arteriosus (PDA)<br>• Total Anomalous Pulmonary Venus Return<br>• SHANK3 involved in myocardial remodeling following myocardial infarction | • (Het) Increased Posterior Wall Thickness at end-diastole (LVPWed)<br>• Increased Anterior Wall Thickness at end-systole (LVAWes) | • PDA can cause PH resulting in an increased wall thickness<br>• ASD can also lead to an increased burden on the ventricular walls resulting in increased wall thickness | (65–68) |
| ***Fmr1*$^{(-/-)}$** | Fragile X Syndrome | • Valvular Heart Diseases - Mitral Valve Prolapse (MVP)<br>• Aortic Root Dilation | • Increased LV Anterior Wall Thickening (LVAWT)<br>• Decreased LV Anterior Wall Thickness at end-diastole (LVAWed) | • MVP causes increased strain on the heart resulting in increased wall thickness<br>• (even mild) MVP causes increased LVEDD | (5,75,93) |

| | | | | | |
|---|---|---|---|---|---|
| | | • Congenital Heart Disease present in <10% of cases | • Increased LV Chamber Diameter in end-diastole (LVEDD) | • LVEDD changes more dramatically than LVESD in less severe stages (compensated normal LV function) | |
| | | • Autonomic Dysfunction | | • Autonomic dysfunction would affect the LVAWT. | |
| | | • Hypertension | | | |
| | | • Cardiac Arrhythmias | | | |
| | | • Arterial Dissection | | | |
| | | • Aneurysm | | | |
| ***Ardi1b***$^{(+/-)}$ | Coffin-Siris Syndrome | • Atrial Septal Defect (ASD) | • Decreased Aorta Diameter (AoD) | • Decreased AoD matches arterial stenosis seen clinically | (4,5) |
| | | • Ventricular Septal Defect (VSD) | • Decreased LV chamber diameter in end-systole and end-diastole (LVESD, LVEDD) | • Arterial stenosis can lead to LV hypertrophy to compensate for increased afterload | |
| | | • Mitral Regurgitation | | • ASD can lead to decrease of LV diastolic area. Here this could be manifested through a diameter decrease. | |
| | | • Patent Ductus Arteriosus | | • VSD can lead to decrease of LV diastolic area. Here this could be manifested through a diameter increase. | |
| | | • Pulmonary Stenosis | | • It's important to note that RV dysfunction | |

| | | | | | |
|---|---|---|---|---|---|
| | | • Arterial Stenosis<br>• Dextrocardia<br>• ARID1B associated with Congenital Heart Disease; present in 20-44% of cases | | can also lead to LV dysfunction. | |
| **16p11.2 (deletion)** | - | • Tetralogy of Fallot (ToF) and associated anomalies<br>• Aortic valve abnormalities (bicuspid valve, stenosis)<br>• Pulmonary atresia<br>• 16p11.2 harbors genes linked to aortic valve development | • Deceased LV Anterior Wall Thickness at end-diastole (LVAWed)<br>• Increased LV Anterior Wall Thickening (LVAWT)<br>• Increased Fractional Shortening<br>• Increased Heart Rate (HR)<br>• Decreased LV Chamber Diameter in end-systole and end-diastole (LVESD, LVEDD) | • Aortic valve dysfunction can result in increased afterload. Then LV remodeling (smaller chamber, increased wall thickness) as well as increased HR are necessary to compensate<br>• Main symptom of ToF is PAH. PAH can result in RV hypertrophy which in turn results in reduction in LVESD<br>• Increase in HR matches ToF symptom of HR irregularities | (5,76,77) |

| | | | | | |
|---|---|---|---|---|---|
| ***Chd8***$^{(+/-)}$ | - | • Involved in cardiac development<br>• Proposed as new AKAP (A-kinase anchoring protein). AKAP dysfunction in the heart linked to hypertrophy, arrythmias and heart failure.<br>• Cardiac defects of CHARGE syndrome through association to CHD7 function.<br>• Tetralogy of Fallot (ToF), Patent Ductus Arteriosus, double outlet RV with AV canal, Ventricular Septal Defect, Atrial Septal Defect (w/ or w/o cleft mitral valve) | • Increased Posterior Wall Thickness at end-systole (LVPWes) and end-diastole (LVPWed). | • Decreased HR may be linked to arrythmias associated with disrupted AKAP function.<br>• CHD8 expressed during both embryonic and postnatal stages and involved in cardiac development from the mesoderm stages; found in myocytes and fibroblasts. All observed changes could have resulted from a developmental dysfunction.<br>• ToF alone can result in HR abnormalities and LV remodeling | (30,81,82,84–86) |
| ***Sgsh***$^{(+/-)}$<br>***Sgsh***$^{(-/-)}$ | Sanfilippo Syndrome A | • No consistent overt cardiac abnormalities | • (Het) Increased E/A | • Diastolic dysfunction can be | (87–89) |

| | | | | | |
|---|---|---|---|---|---|
| | | | | reflected through changes in the E/A ratio; inconsistency with reduced E/A ratio reported. | |
| | | • Mitral Valvulopathy | • (Hom) Increased Fractional Shortening (FS) | • Changes in fractional shortening can reflect the changes in shape of LV seen clinically | |
| | | • Ventricular Septal Defect | • (Hom) Increase in LV Anterior Wall Thickening (LVAWT) | • LV dysfunction can be reflected in the change of LVAWT. | |
| | | • Subclinical Systolic Dysfunction<br>• Early LV Dysfunction (increased stress with normal Ejection Fraction)<br>• LV Diastolic Dysfunction<br>• Reduced Ejection Fraction<br>• Congestive Cardiomyopathy<br>• LV Enlargement | | | |
| ***Vps13b$^{(+/-)}$*** | Cohen Syndrome | • Decreased LV function with advancing age<br>• Normal heart anatomy, Aorta Diameter and LV wall thickness<br>• Essential Hypertension | • Decreased LV Anterior Wall Thickness at end-diastole (LVAWed) | Similarly to clinical findings, hardly any abnormalities were reported. | (90–92) |

| | | |
|---|---|---|
| | | • Pulmonary Hypertension (PH)<br>• Cardiac Systolic Murmurs<br>• Inconsistent valvular and vascular defects, and myocardial abnormalities (as detected with EEG) |

*Table 5: Documentation of the clinically reported cardiac phenotypes (column 3) for the ASD-related genetic models used in the current study (column 1) along with the cardiac abnormalities observed in the current study (column 4). Distinction between heterozygous and homozygous genotype (in column 4) made ony in cases where both genotypes were used. Note there is no implied association between the entries of columns 3 and 4 of the same row; there is no relation between the sequence of listed observations between the two columns. Column 2 gives the syndrome associated with each genetic knockout. Column 5 gives the potential association between the phenotypes presented in columns 3 and 4. LV=left ventricular, Het=heterozygous for given gene, Hom= heterozygous for given gene.*

<u>Total Correlation Calculation</u>

Total correlation $C(X_1, X_2, \ldots, X_N)$ is the difference between the sum of the information entropy of each variable $\{X_i\}$ in a set $\{X_1, X_2, \ldots, X_N\}$ and the joint entropy of the variable set $\{X_1, X_2, \ldots, X_N\}$ (equation 4). $C(X_1, X_2, \ldots, X_N)$ represents the amount of information shared between the components of a dataset, or alternatively the redundancy in a dataset.

$$C(X_1, X_2, \ldots, X_N) = \left[\sum_{i=1}^{N} H(X_i)\right] - H(X_1, X_2, \ldots, X_N)$$

*Equation 4: Total correlation $C(X_1, X_2, \ldots, X_N)$ for a dataset $\{X_1, X_2, \ldots, X_N\}$. $H(X_i)$ is the information entropy of variable $X_i$ and $H(X_1, X_2, \ldots, X_N)$ the joint entropy of the variable set $\{X_1, X_2, \ldots, X_N\}$.*

Each dataset used in our analysis was a variable set and each column was a variable. To calculate the entropy of a continuous variable or variable set (differential entropy) the probability density function is needed. For our data, the method of moments was employed to estimate the probability density function per variable and variable set. Following inspection of the data, the normal distribution (univariate and multivariate respectively) was chosen as the theoretical distribution. The first (mean) and second moment (variance $\sigma_i$ per variable $X_i$ and covariance Σ per variable set $\{X_1, X_2, \ldots, X_N\}$) were estimated.

For a normal probability density function (PDF), the differential entropy is given by:

$$(A) \qquad H(X_i) \sim ln\left(\sigma_i \cdot \sqrt{2\pi e}\right) \qquad \textit{for univariate normal PDF}$$

$$(B) \qquad H(X_1, X_2, \ldots, X_N) \sim \frac{1}{2} \cdot ln\{(2\pi e)^N \cdot \det(\Sigma)\} \qquad \textit{for multivariate normal PDF}$$

*Equation 5: Differential entropy for (A) a variable $X_i$ with a univariate normal probability density function and variance $\sigma_i$; (B) a variable set $\{X_1, X_2, \ldots, X_N\}$ with a multivariate normal probability density function with covariance matrix Σ and $\det(\Sigma)$ the determinant of Σ.*
